# OmniReprodubileCellAnalysis: a comprehensive toolbox for the analysis of cellular biology data

**DOI:** 10.1101/2023.11.07.565961

**Authors:** Tortarolo Dora, Pernice Simone, Fabiana Clapero, Donatella Valdembri, Guido Serini, Federica Riccardo, Lidia Tarone, Chiara Enrico Bena, Carla Bosia, Sandro Gepiro Contaldo, Beccuti Marco, Marzio Pennisi, Cordero Francesca

## Abstract

Open science and reproducibility are two key pillars of modern scientific research. Open science is making scientific research and data accessible and transparent to the broader scientific community and the public. Reproducibility, on the other hand, is the ability to replicate and confirm research results by following the same methods and procedures. Reproducibility is thus crucial because it ensures the reliability and validity of scientific findings. The relationship between open science and reproducibility is intertwined; indeed open science practices, such as sharing raw data, detailed methodologies, and code, greatly facilitate the reproducibility of research. In recent years, concerns about the reproducibility of scientific research have gained prominence, and indeed scientists still lament the lack of details in the methods sections of published papers and the unavailability of raw data from the authors. To assist cellular biologists and immunologists and to promote a more transparent, open and reproducible research practice, we developed *OmniReproducibleCellAnalysis* (*ORCA*), a new Shiny Application based in R, for the semi-automated analysis of Western Blot (WB), Reverse Transcription-quantitative PCR (RT-qPCR), Enzyme-Linked ImmunoSorbent Assay (ELISA), Endocytosis and Cytotoxicity experiments. *ORCA* is open-source and approachable by scientists without advanced R language knowledge. Our application automatically compiles a report containing the finalized data analysis and all its preliminary and intermediate steps, ensuring data analysis standardization and reproducibility. Furthermore, *ORCA* allows to upload raw data and results directly on the data repository Harvard Dataverse, a valuable tool for promoting transparency and data accessibility in scientific research. By employing *ORCA*, scientists will cut down analysis time and human-dependent errors, while taking a step towards a research practice compliant with Open Science and FAIR principle.

## Introduction

In recent years, the scientific community has been shaken by the notion that most of the scientific results published even in foundational research papers can not be reproduced. In the field of immunology, reproducibility is not well-defined for most of the immunological parameters that are often involved in complex mechanisms [1]. Similarly, in the field of cancer research, studies show that only about 20-25% [2] or 11% [3] of published studies can be validated or reproduced. The results of independent surveys investigating researchers’ experience of experiment reproducibility [4,5] convene that more than 70% of participants have failed to replicate other scientists’ data.

While the debate is still open on whether the lack of reproducibility is due to the intrinsic variability of living organisms and of the reagents employed to study them [6], most participants blame incomplete specification of original protocol to accurately guide the replication attempt [5] and unavailability of raw data from the original laboratory [4]. Indeed, most scientific papers fail to specify the peculiar details of the protocols employed: they may generically refer to the methodological sections from other papers, or provide vague and subjective directions such as “stir until yellow”. Often scientific papers leave out altogether how the analysis of the raw data is carried out, making it impossible to fully reproduce results, even with the raw data at hand. Exemplary is the analysis of Western Blot (WB) using an open access software such as ImageJ, where densitometric curves are manually cut by the user to distinguish between signal and noise. Each cut is irreproducible since it is user-dependent and usually not recorded for future validation.

Additionally, the raw data is often not available, even upon request to the authors [7]. At the same time, research results are mostly displayed as representative figures or visualizations of finalized datasets that have undergone extensive manipulation and do not reveal the original data [8]. These published results are not suitable for data mining, reuse, validation, and reproduction. In light of the growing awareness of reproducibility concerns, stakeholders such as funders, publishers, and policymakers have come together to promote Open Science practices, to delineate strategies to carry out a more transparent research practice at all levels of scientific endeavors [9–11], and to implement the FAIRness principles of data Findability, Accessibility, Interoperability, and Reusability, for effective management and sharing of data holdings [12].

Yet, in practice, the current software available for the analysis of the most common cellular biology and immunological experiments lacks the tools and characteristics to easily and straightforwardly put into effect the aforementioned guidelines, leaving to the scientist the tedious labor of manually reporting thresholds, variables, and bits of scripts.

Moreover, to the best of our knowledge, comprehensive software where the scientist can perform all of the above analyses is still missing. Laboratories often opt to perform data analysis on the proprietary software provided with the instrument employed to carry out the experiment. Their licenses are often restrictive and hinder data sharing between collaborators. Additionally, in immunoinformatics, the data-driven computational and statistical approaches are fundamental to describing the dynamics of immunological systems at both cellular and molecular levels. The need to create an interoperable environment to analyze, visualize, and share omni datasets according to the FAIR principle becomes urgent and mandatory.

Addressing this aspect several general-purpose data repositories, such as Harvard Dataverse [13], FigShare [14], DataHub [15], and Zenodo [16], are gaining popularity within the academic research community. They embody FAIR principles by storing, organizing, and uniquely identifying digital resources. Developed at Harvard to solve the problem of data sharing, Dataverse is a trusted platform for open-access data storage, offering researchers a secure and collaborative environment for sharing, preserving, and managing their datasets. Its features include data organization, version control, data citation, and integration with research workflows, making it a valuable tool for promoting transparency and data accessibility in scientific research. However, scientists may view manually uploading laboratory data and results to an online repository as an additional burden on their already heavy workload. Indeed, skipping from one tool to the next to analyze, store, and share data may disincentivize even the more enthusiastic researcher.

However, as suggested by Florian Markowetz in [17], the main beneficiaries of reproducible research practice are the researchers themselves. With this in mind, we developed *ORCA* to support them throughout the scientific discovery process. *ORCA* is a toolbox for the cellular biologist or immunologist to analyze the most common laboratory experiments, including WB, Reverse Transcription-quantitative PCR (RT-qPCR), Cytotoxicity, Enzyme-Linked ImmunoSorbent Assay (ELISA), and Endocytosis experiments. It has been created to comply with Open Access research practices and FAIR principles. On *ORCA*, users can upload raw data, label samples, and experimental conditions, and perform a semi-automated data analysis that cuts time and opportunity for human mistakes. Detailed reports of the data analysis procedure and the resulting outcomes can be automatically downloaded in different formats or published directly on Dataverse. Moreover, *ORCA* provides a streamlined and effective method for automatically fetching datasets and their related details from Dataverse.

## Tool Architecture

*ORCA*, as shown in Fig.1, is a web application created with the open-source Shiny package in R, designed to be user-friendly and accessible to all individuals, including those with minimal or no prior knowledge of the R language. It is freely accessible on GitHub at https://github.com/qBioTurin/ORCA or through the Docker image https://hub.docker.com/repository/docker/qbioturin/orca-shiny-application/general. Note that the use of *ORCA* within a Docker image [18] empowers users to enhance the reproducibility of data analysis.

**Figure 1.**
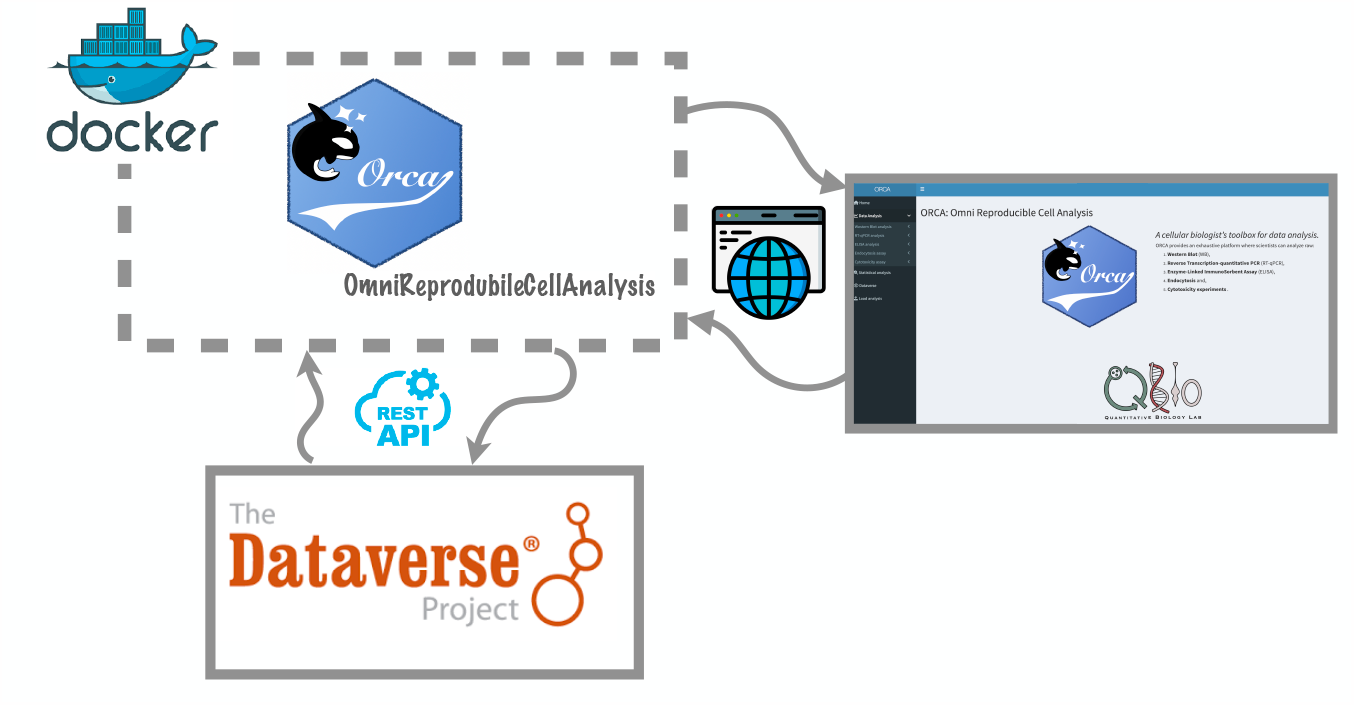
Orca architecture schema.

In detail, *ORCA* is composed of four main modules:

1. Data Analysis
2. Statistical Analysis
3. Load Analysis
4. Dataverse

In the “Data Analysis” module, users can carry out a full data analysis process, from raw data to a finalized figure, for the following experiments: WB, RT-qPCR, ELISA, Endocytosis and Cytotoxicity assays. A thorough description of how *ORCA* works can be found in section Tool Description.

Moreover, the analysis is recorded throughout its entire process. If needed, users can freeze the analysis at any point in time by saving and downloading a .rds file, which can be later uploaded again to *ORCA* to complete the task. Once the analysis is finalized, scientific results and a detailed report of all steps of the analysis can be exported as a .rds and a .xlsx file.

In the “Load Analysis” module, users can upload a .*rds* file either to complete a draft analysis or to perform a statistical study of the completed analysis.

In the “Statistical Analysis” module, users can perform basic statistical analysis on multiple assays, calculating the average, standard deviation, and Student’s t-test of two or more experiments.

In the “Dataverse” module, users can upload results directly on Harvard Dataverse, to deposit and share their data in compliance with the FAIR principles. On Dataverse, users can provide detailed information (as metadata) on how the experiment and the analysis were carried out, and upload the final results. A persistent identifier (e.g. Digital Object Identifier (DOI)) can be assigned to the data once the user decides to publish them from their own account.

## Tool Description

In this section, we will walk the reader through the analysis of WB, RT-qPCR, ELISA, Endocytosis, and Cytotoxicity assays. We will explain how to perform final statistical analysis to compare multiple results of a specific analysis and how to upload results on Dataverse. Each type of analysis will be performed using a toy example based on real data, either unpublished or already printed.

### 0.1 Data Analysis

By clicking on the “Data Analysis” section of the menu on the left side of the interface, users can select the type of experiment of interest. Let us note that a full .pdf report of the analysis can be downloaded by clicking on the purple button on the top right-hand side of the screen.

#### Western Blot analysis

Herein, we will analyze a WB experiment on primary human umbelical arthery endothelial cells (HUAECs) [19] treated with 200 pmol of siGENOME Non-Targeting siRNA #1 (as siCTL) or siGENOME SMART pools siRNA oligonucleotides for PPFIA1, using Oligofectamine Transfection Reagent (Life Technologies), according to the manufacturer’s protocol. The day before oligofection, cells were seeded in 6-well dishes at a concentration of 12 *∗* 10^4^ per well. HUAECs were either treated once (at 0h) and lysed after 12 or 24h, or twice (at 24h) and lysed after 12 or 24h.

The user can upload the WB image by selecting the “Upload Image” section from the drop-down menu. Here, it is possible to browse for the WB image of interest. The image should be a tiff file. By clicking on the “Load” button, the image is uploaded and the user is automatically re-directed to the “Protein Bands” section. On this page, the user can select the protein bands of interest by drawing a rectangle over the area and clicking on “Select Protein Band” (see Fig. 2, A1). The coordinates of the rectangle are stored on the left-hand side of the window (“Protein Band Selection Coordinates” box, see Fig. 2, A2), and they are recorded in the analysis report to ensure full reproducibility. By dragging the rectangle, it is possible to select multiple protein bands. Once all protein bands of interest have been selected, the user can click on the “Generate Plots” button to be re-directed to the “Profile Plots” section to analyze the densitometric curves (Fig. 2, A3). In the “Profile Plots” section, the scientist can remove background noise by truncating the densitometric peak. Users can perform a vertical and/or horizontal truncation by adjusting the sliders in the “Truncation” box. Each cut and Area Under the Curve (AUC) are stored in the lower left-hand side and recorded in the analysis report, which the user can download as a .rds or .xlsx file by clicking on the “Download the analysis” button. In the “Quantification” section, the user should upload two .rds files: (1) containing the analysis of the WB of the protein of interest and (2) the analysis of the WB containing a normalizer (or housekeeping gene). The .rds contains for each WB lane all of the intermediate and final AUCs with respective truncation coordinates. For each sample, the user should select the final truncation by clicking on the correct row (see Fig. 2, A4). In the “Relative Density” box, the user can normalize the samples from the protein of interest WB on its control sample. Users must select the control sample to use for the relative normalization from the drop-down menu. In the “Adjusted Relative Density” box, the AUC from the protein of interest is normalized on the AUC of the normalizer WB. Since the two WB share sample order, the Adjusted Relative Density is automatically calculated. A graph displays the Adjusted Relative Density for all samples compared to the control sample baseline (see Fig. 2, A5). By clicking on the “Download the analysis” button, the user can download an .rds containing all steps of analysis. Results can also be downloaded as a .xlsx file, by clicking on the “Download xlsx” button.

**Figure 2.**
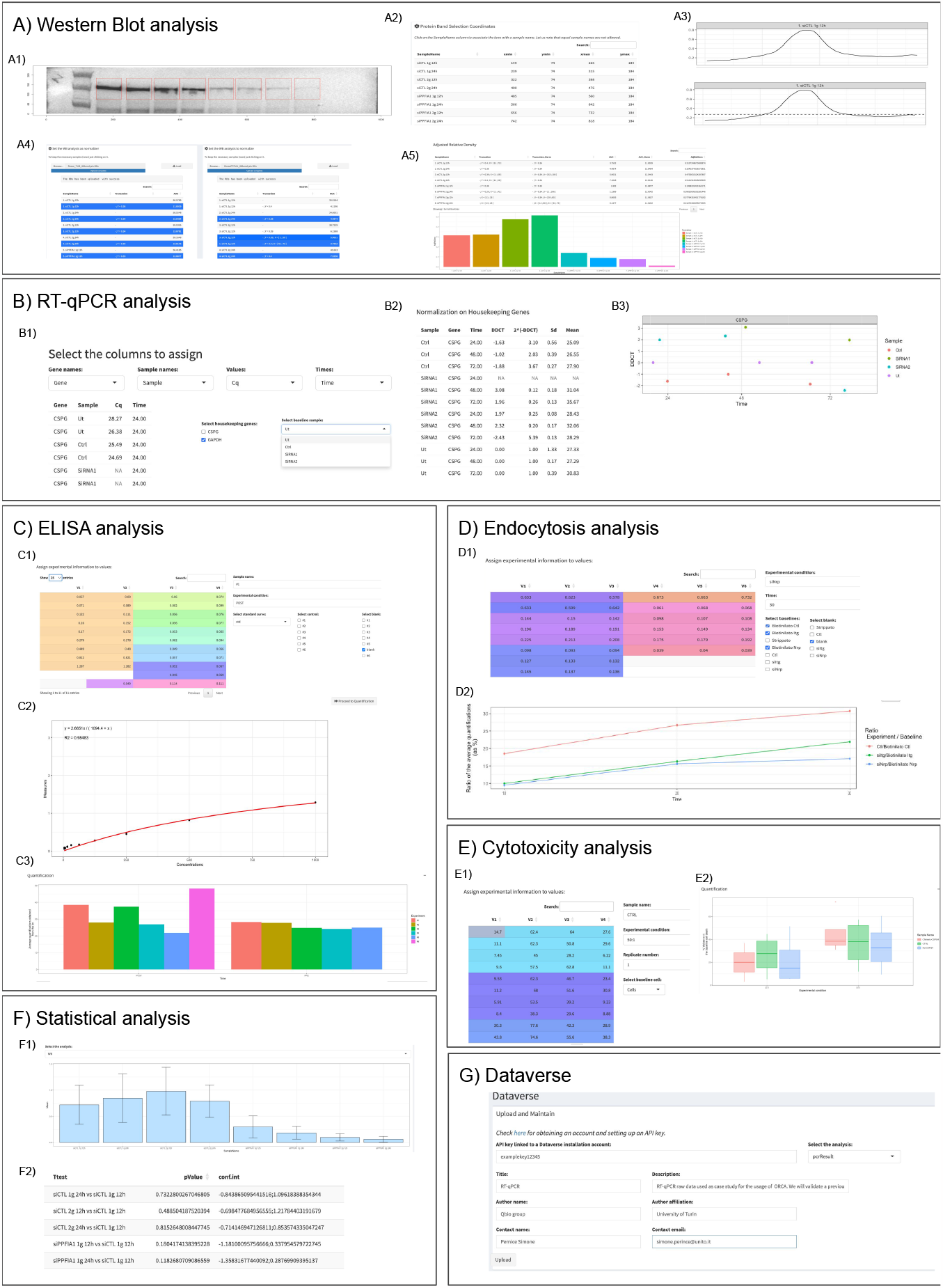
Summary of all the steps of analysis for each WB (A), RT-qPCR (B), ELISA (C), Endocytosis (D), Cytotoxicity (E), Statistical Analysis (F), and Dataverse (G) sections in *ORCA*.

#### RT-qPCR analysis

To illustrate the analysis of RT-qPCR raw data using *ORCA*, we will validate a previously published experiment [20]. This assay investigates the functional consequences of silencing the Chondroitin Sulfate Proteoglycan (CSPG4) gene with two different anti-CSPG4-siRNA in the human U2OS osteosarcoma cell line. For details on how the experiment was carried out, we refer to [20]. In the “Upload Data” section, users can upload a .xlsx file containing a dataset encoded as a rectangular data structure in which the columns correspond to the variables describing each entry (e.g. samples, gene targets, measured threshold Cycle (Ct), and time). In the “Experimental Setup” box, the scientist can match the column with the appropriate description (see Fig, 2, B1). *ORCA* will suggest selecting the housekeeping genes and request to select which sample is the control (Fig. 2, B1). By clicking on “Proceed to quantification”, *ORCA* will automatically calculate the relative expression of target genes using the 2^(*−*ΔΔ*ct*)^ method [21]. In particular, grouping the data w.r.t. the samples, gene targets, and the time values, the average measured Ct is calculated and then, once the housekeeping gene average Ct is subtracted, the Δ*ct* is defined. Successively,*ORCA*calculates the ΔΔ*ct* by subtracting the control sample Δ*ct* from all samples. Thus, *ORCA* will provide as output the normalized 2^(*−*ΔΔ*ct*)^ values on housekeeping genes. The output is visualized as a table (Fig. 2, B2) and as graphs (Fig. 2, B3). By clicking on the “Download the analysis” button, the user can download a .rds containing all steps of analysis. Results can also be downloaded as a .xlsx file, by clicking on the “Download xlsx” button.

#### ELISA analysis

To illustrate the analysis of ELISA raw data using *ORCA*, we will validate a previously published experiment [20]. This experiment investigates the release of canine Interferon *γ* (IFN-*γ*) by Peripheral Blood Mononuclear Cells (PBMC) collected before (Pre-Vax) and after (Post-Vax) the IV immunization with a chimeric-CSPG4 coding plasmid, in the presence of dog (Do)-CSPG4 peptides (PEP). For details on how the experiment was carried out, we refer to [20].

In the “Upload Data” section, users can upload a .xlsx file containing a dataset encoded as a rectangular data structure in which the columns correspond to the variables describing each entry (e.g. standard curve and sample measurements). When the file has been uploaded successfully, *ORCA* automatically loads the raw data in the “Assign experimental information to values” box. Here, the user can click on each value from the raw data and assign the “Sample Name” and “Experimental Condition” labels (see Fig. 2, C1). When written once, *ORCA* the name is stored and can be added simply by clicking on the option from the drop-down menu. Let us note that if multiple values are associated with the same sample name and experimental condition then the average of these values will be considered through the next steps. The scientist can identify among all values which belong to the standard curve, by selecting the appropriate “Sample Name” label from “Select Standard Curve” in the drop-down menu (Fig. 2,C1). They can classify the control and blank samples from the “select control” and “select blank” options (Fig. 2, C1). By clicking on “Proceed to Quantification”, it is possible to automatically reach the “Quantification” section, where the user can i) add to the standard curve measurements the corresponding concentration values, and ii) select between two type of regression models, linear or hyperbola, to extrapolate sample quantification from the standard curve. Specifically, the plot depicting the data and model selected is shown together with the linear model equation *y* = *m ∗ x* + *q* or the hyperbola model *y* = *Bmax ∗ x/*(*Kd* + *x*), with the corresponding parameters estimated by fitting the model with the standard curve data and the coefficient of determination, denoted *R*^2^. In Fig. 2 C2, the hyperbola regression model obtained from the selected standard curve is shown. The estimated model is defined by the equation *y* = 2.6651 *∗ x/*(1094.4 + *x*) with *R*^2^ = 0.98483.

Successively, the estimate model is exploited to quantify all the sample average measurements subtracted by the average blank value. Finally, a bar plot reporting the resulting quantification values grouped by the experimental condition is automatically generated, as in Fig. 2, C3.

By clicking on the “Download the analysis” button, the user can download a .rds containing all steps of analysis. Results can also be downloaded as a .xlsx file, by clicking on the “Download xlsx” button.

#### Endocytosis assay

An endocytosis assay evaluates the percentage of surface proteins that are endocytosed in a given time frame. In our example, we perform a time-course analysis of the relative amounts of endocytosed Kinase insert Domain Receptor (KDR), also known as Vascular Endothelial Growth Factor Receptor 2 (VEGFR), in HUAECs. HUAECs were treated twice (at 0 and 24h) with 200 pmol of siGENOME Non-Targeting siRNA #1 (as siCTL) or siGENOME SMART pools siRNA oligonucleotides for Nrp1 and for ItgB1, using Oligofectamine Transfection Reagent (12252-011, Life Technologies), according to the manufacturer’s protocol. The integrin endocytosis assay and capture ELISA assays were performed 24 hours after the second oligofection, as described in [19].

In the “Upload Data” section, users can upload a .xlsx file containing a dataset encoded as a rectangular data structure in which the columns correspond to the variables describing each entry (e.g. replicates of sample values). Once loaded, the user can assign a label, by clicking on the value and completing writing out the name of the experimental condition in the “Experimental Condition” drop-down menu, and the time interval of analysis in the “Time” drop-down menu (see Fig. 2, D1). Let us note that if multiple values are associated with the same experimental condition and time then the average of these values will be considered through the next steps. When written once, *ORCA* the name is stored and can be added simply by clicking on the option from the drop-down menu. The user can classify the baselines and blank samples from the “Select baselines” and “Select blank” options (Fig. 2, D1). By clicking on “Proceed to Quantification”, after each experimental condition is associated with the corresponding baseline sample and blank, the ratio between each average experimental condition and the corresponding baseline is automatically returned grouped by the time and experimental condition variables as shown in Fig. 2, D2.

By clicking on the “Download the analysis” button, the user can download a .rds containing all steps of analysis. Results can also be downloaded as a .xlsx file, by clicking on the “Download xlsx” button.

#### Cytotoxicity assay

To illustrate the analysis of a cytotoxicity assay using *ORCA*, we will validate a previously published experiment [20]. A cytotoxicity assay was performed with healthy donor PBMC recovered after 7 days of co-culture with autologous dendritic cells transfected with either the empty vector, the human (Hu)-CSPG4, or the chimeric-CSPG4 plasmids. Pre-activated PBMC were incubated with CFSE-labeled CSPG4+ U2OS human Osteosarcoma (OSA) cells, at different effector:target (E:T) ratios. For details on how the experiment was carried out, we refer to [20]. In the “Upload Data” section, users can upload a .xlsx file containing a dataset encoded as a rectangular data structure in which the columns correspond to the variables describing each entry (e.g. replicates of sample values). Once loaded, the user can assign a label, by clicking on the value and completing writing out the name of the sample in the “Sample name” drop-down-menu, the experimental condition in the “Experimental Condition” drop-down menu, the experimental replicate in the “Replicate number” drop-down menu, and the samples belonging to measures of basal cell death in the “Select baseline cell” drop-down menu (see Fig. 2, E1). When written once, *ORCA* the name is stored and can be added simply by clicking on the option from the drop-down menu. By clicking on “Proceed to quantification”, *ORCA* automatically calculates the percentage of cell death with respect to the baseline and plots the results in a graph (see Fig. 2, E2). *ORCA* calculates the cytotoxicity using the well-established formula

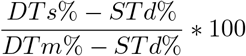

where *STd*% represents the percentage of spontaneously dead targets, *DTs*% the percentage of dead targets in sample, and *DTm*% the maximum percentage of dead target. By clicking on the “Download the analysis” button, the user can download a .rds containing all steps of analysis. Results can also be downloaded as a .xlsx file, by clicking on the “Download xlsx” button.

### 0.2 Statistical analysis

In the “Statistical Analysis” module, the user can upload multiple .rds files containing finalized analyses of the same type (e.g., WB). In the “Comparison Analysis” box, they can select the analysis of interest from a drop-down menu which is automatically updated with the analysis name stored in the rds files. An example considering three WB quantification analysis is depicted in Fig. 2, F. Specifically, the average and standard deviation considering the quantification results of the selected analysis are calculated and plotted (Fig. 2, F1). Finally, granted that the analysis are of the same type, and defined by the same sample names, experimental conditions and time values (depending on the analysis), a statistical t-test is used to compare the means of two groups generated by all the possible combinations between two different sample names (in case of the WB, RT-qPCR, ELISA, and Cytotoxicity analysis) or experimental conditions (in case of the Endocytosis analysis). Fig. 2, F2 shows a subset of combinations among all the eight sample names characterising the WB analysis.

### 0.3 Dataverse

This module was designed for researchers to effortlessly upload and share their research datasets on the Harvard Dataverse platform. It harnesses the Dataverse API, offering a seamless and intuitive process for data contributors implemented into an ad-hoc developed Python library, encapsulated into the docker image *qbioturin/orca-upload-dataverse*. Indeed, the module guides users through dataset creation, making it easy to provide comprehensive metadata, such as titles, authors, descriptions, keywords, and licensing information. Thus, researchers can efficiently upload their raw data files and, if desired, multiple versions of their analysis outputs specifying their visibility settings. Finally, once succesfully published, a DOI will be assigned to each dataset, simplifying data citation and referencing. Indeed, the authors should register to https://demo.dataverse.org/ to have an account and to set up an API key that will be exploited to upload the analysis on their own account. Specifically, for each type of analysis two files will be uploaded: the excel and the .rds files storing the raw data, and all the analysis result obtained through *ORCA*.

An example of the Dataverse section filled to upload a RT-qPCR analysis is depicted in Fig. 2, G.

## Conclusions

In recent years, the volume of experimental data generated has continuously increased, requiring the establishment of comprehensive computational environments to enable easy and intuitive data analysis, and to foster a more transparent, open, and reproducible research practice.

To the best of our knowledge, *ORCA* is the first comprehensive free environment able to perform data analysis and store data following the FAIR principles. *ORCA* is a powerful and fruitful web toolbox to help the user in the research practice by providing a semi-automatized data analysis of five types of experiments WB, RTqPCR, ELISA, Endocytosis, and Cytotoxicity assays. Furthermore, it offers detailed reports of the data analysis process, and the resulting outcomes can be automatically retrieved in various formats or uploaded directly on a Dataverse instance.

In the future, the following three crucial aspects will be addressed to further enhance the usability of *ORCA*. Firstly, to facilitate the scientist in carrying out a reproducible research practice, we will extend *ORCA*’s functionalities to accommodate a Section where (i), before embarking on their scientific investigation, they can delineate their study design in a Registered Report [22], and (ii), throughout their discovery process, they can keep an Electronic Laboratory Notebook (ELN) to digitally document how the experiment was conducted, report details of data and link them to the results. A fairly recent practice, publishing a Registered Report rewards best practices by conducting a peer review prior to data collection and ensuring a publication in peer-reviewed journals, if the authors follow through with the registered methodology, irrespectively of research outcomes [8].

Secondly, for each type of experiment, we will create a specific *Analytical Chamber* to analyze and visualize the results. In each *Analytical Chamber* a unique set of analytical methods will be provided to perform both the descriptive and inference statistics specifically for the experiment selected. In this manner, the user should be confident in using the appropriate methods to explore the data collected.

Finally, to give the user the opportunity to globally envision the phenomenon under study, the use of computational models helps improve the understanding of the biological/clinical contexts. One of the major criticalities in employing computational models is setting the parameters. We are strongly confident that several model parameters could be extrapolated just by observing the experiments from a new perspective. An environment such as *ORCA* gives the opportunity to perform an interactive data exploration of several types of quantitative experimental data, which can be useful in defining an in-silico model to test new biological hypotheses or new theories.

## Acknowledgements

This work was partially supported by CINI infoLife laborstory through the “La resilienza del Mar Mediterraneo al cambiamento climatico direzionale: sviluppo di modelli sperimentali e di rischio” project funded by Ministero della Salute - Ricerca Corrente 2022. M.P. would like to thank Mimesis S.r.l. for financial support.

## Glossary

AUC: Area Under the Curve. 5
CSPG4: Chondroitin Sulfate Proteoglycan. 5–7
Ct: threshold Cycle. 6
DOI: Digital Object Identifier. 4, 9
ELISA: Enzyme-Linked ImmunoSorbent Assay. 3, 4, 9, 12
ELN: Electronic Laboratory Notebook. 9
HUAECs: human umbelical arthery endothelial cells. 5, 7
IFN-γ: Interferon γ. 6
KDR: Kinase insert Domain Receptor. 7
OSA: Osteosarcoma. 7
PBMC: Peripheral Blood Mononuclear Cells. 6, 7
RT-qPCR: Reverse Transcription-quantitative PCR. 3, 4, 9, 12
VEGFR: Vascular Endothelial Growth Factor Receptor 2. 7
WB: Western Blot. 2–5, 8, 9, 12

## Notes

### Competing Interest Statement

The authors have declared no competing interest.

